# DeepGWAS: Enhance GWAS Signals for Neuropsychiatric Disorders via Deep Neural Network

**DOI:** 10.1101/2022.12.20.521277

**Authors:** Jia Wen, Gang Li, Jiawen Chen, Quan Sun, Weifang Liu, Wyliena Guan, Boqiao Lai, Haibo Zhou, Jin P Szatkiewicz, Xin He, Patrick F. Sullivan, Yun Li

**Affiliations:** Department of Genetics, University of North Carolina at Chapel Hill, Chapel Hill, NC, 27599, USA; Department of Statistics and Operations Research, University of North Carolina at Chapel Hill, Chapel Hill, NC, 27599, USA; Department of Genome Sciences, University of Washington, William H. Foege Hall, 3720 15th Ave NE, Seattle, WA, 98195, USA; eScience Institute, University of Washington, WRF Data Science Studio, UW Physics/Astronomy Tower, 6th Floor, Campus Box 351570, 3910 15th Ave NE, Seattle, WA 98195, USA; Department of Biostatistics, University of North Carolina at Chapel Hill, Chapel Hill, NC, 27599, USA; Toyota Technological Institute at Chicago, Chicago, Illinois, United States of America; Department of Psychiatry, University of North Carolina at Chapel Hill, Chapel Hill, NC, 27599, USA; Department of Human Genetics, University of Chicago, Chicago, Illinois, United States of America; Department of Medical Epidemiology and Biostatistics, Karolinska Institutet, Stockholm, Sweden, 17177

**Author notes:** These authors contributed equally as corresponding authors. These authors contributed equally as first authors (Jia Wen, Gang Li).

## Abstract

Genetic dissection of neuropsychiatric disorders can potentially reveal novel therapeutic targets. While genome-wide association studies (GWAS) have tremendously advanced our understanding, we approach a sample size bottleneck (i.e., the number of cases needed to identify >90% of all loci is impractical). Therefore, computationally enhancing GWAS on existing samples may be particularly valuable. Here, we describe DeepGWAS, a deep neural network-based method to enhance GWAS by integrating GWAS results with linkage disequilibrium and brain-related functional annotations. DeepGWAS enhanced schizophrenia (SCZ) loci by ∼3X when applied to the largest European GWAS, and 21.3% enhanced loci were validated by the latest multi-ancestry GWAS. Importantly, DeepGWAS models can be transferred to other neuropsychiatric disorders. Transferring SCZ-trained models to Alzheimer’s disease and major depressive disorder, we observed 1.3-17.6X detected loci compared to standard GWAS, among which 27-40% were validated by other GWAS studies. We anticipate DeepGWAS to be a powerful tool in GWAS studies.

---

Neuropsychiatric disorders carry high public health burden including tremendous morbidity, mortality, lessened quality of life, and financial costs^1,2^. For example, schizophrenia (SCZ) is a highly heritable and debilitating psychiatric disorder affecting about 0.28% of the global population and is associated with high morbidity, mortality, as well as personal and public health costs^3^. During the past 15 years, GWAS have greatly advanced our understanding of the genetic basis underlying these disorders^4,5^. For example, SCZ started with 1 locus reaching genome-wide significance in a GWAS with 3,322 cases in 2009^6^ to 287 loci in the most recent meta-analysis^5^ of ∼75K cases. The tremendous advancement is largely attributable to increased sample size, which is of undisputed value in GWAS for many complex diseases^7^. However, increasing sample size by another order of magnitude in GWAS becomes increasingly challenging, particularly for SCZ and other neuropsychiatric disorders. Therefore, enhanced GWAS on existing samples via computational approaches would be particularly valuable for genetic dissection of neuropsychiatric disorders.

Standard GWAS associates genotypes with phenotypes usually assumes that all variants are *a priori* equally likely to be associated^7^. This assumption was initially proper as priors were either unavailable or debated. However, we have now accumulated rich genomic and epigenomic evidence and the continuation of this assumption may represent a tremendous, missed opportunity to leverage and integrate standard GWAS results with functional annotations to effectively up-weight variants that are more likely to play functional roles. For example, GWAS variants have been reported to enrich in regulatory regions^8,9^, and explain a larger than expected amount of disease and trait heritability^10,11^. Therefore, leveraging functional annotations could enhance statistical power to identify causal variants. Researchers have employed similar integration ideas for related purposes, including phenotypic prediction, gene-gene interaction detection and post-GWAS prioritization of genetic variants and their target genes^12-16^ but not for GWAS *per se*.

Here, we apply machine learning to integrate summary statistics from standard GWAS with functional annotation information for enhancing GWAS findings. Specifically, we develop DeepGWAS, a 14-layer deep neural network to enhance GWAS signals without increasing sample size. The input predictors include GWAS summary statistics, linkage disequilibrium (LD) information, and brain related functional annotations. We first trained our DeepGWAS model with SCZ GWAS summary statistics, finding that our DeepGWAS model outperformed other state-of-the-art machine learning and traditional statistical methods, including XGBoost^17^ and logistic regression. Encouraged by these results, we further transferred our DeepGWAS model trained on SCZ to enhance GWAS for two other neuropsychiatric diseases.

## Results

### Overview of the DeepGWAS model

DeepGWAS infers the probability of a variant associated with the phenotype of interest by modeling a vector of 33 input features in a 14-layer fully connected deep neural network (**1** and **Online Methods**). In DeepGWAS models, genetic variants are observations, and for each observation, the input features include GWAS summary statistics, basic population genetics statistics, and brain-related functional annotations (**Online Methods**). Training a DeepGWAS model entails label information (i.e., the binary label indicating whether a variant is associated with the phenotype of interest). In reality, gold-standard true labels typically do not exist.

Therefore, we recommend training a DeepGWAS model using results from two GWA studies where the smaller and less powerful GWA study provides the input features while the more powerful one provides label information. The trained DeepGWAS models can be applied to enhance the more powerful GWAS (which only provides input features), enhance another GWAS for the same phenotype, or enhance GWAS for another brain-related disease (for examples, see **“DeepGWAS enhancement reveals hundreds of novel SCZ loci”, “Enhancing AD GWAS” and “Enhancing MDD GWAS*”*** sections below).

### Systematic evaluation of DeepGWAS using SCZ GWAS data

We first compared the ability of DeepGWAS to enhance GWAS signals with two alternative methods, logistic regression and XGBoost^17^. The former is a classic statistical method, and the latter is a widely-used machine learning method. Results show that DeepGWAS achieves the best performance at both variant and locus-level (**Fig. 2**). Using GWAS summary statistics from 64 SCZ GWAS cohorts^5^, we were able to design careful experiments (**Table S1** and **Table S2**) for systematic evaluation. Since neural networks are prone to a trivial solution due to a highly imbalanced data such as GWAS summary statistics, DeepGWAS adopts an under-sampling strategy for non-significant variants when selecting a subset of variants for training (detailed in “**Under-sampling insignificant variants for model training*”*** in **Supplementary Materials**). For logistic regression and XGboost, we consider models trained on the full sample of variants as well as models trained on the subset sample of variants as input into the DeepGWAS model (indicated by “_subset”). Each of the five models takes the same 33 features as input. With the default prediction probability threshold value of 0.5, DeepGWAS achieved first place (at variant level) and second place (at locus level) for capturing true positives and had an overall best and second best F1 score balancing sensitivity and specificity at the variant level and locus level, respectively (**Fig. 2**). For example, at the locus level, the F1 score of XGBoost (0.07) was less than half that of DeepGWAS (0.16). Although logistic regression applied to all variants had the highest F1 score, DeepGWAS approximately doubled the power (TPR: 0.28 vs 0.55). Thus DeepGWAS provided the best balance between power and overall performance (**Fig. 2**). At the variant level, DeepGWAS (red curve) outperforms all other models and is the clear winner in terms of power (TPR) with a range of 40-60%, the only range where methods have reasonable power and acceptable false positive control (**Fig. 2**).

**Fig. 1.**
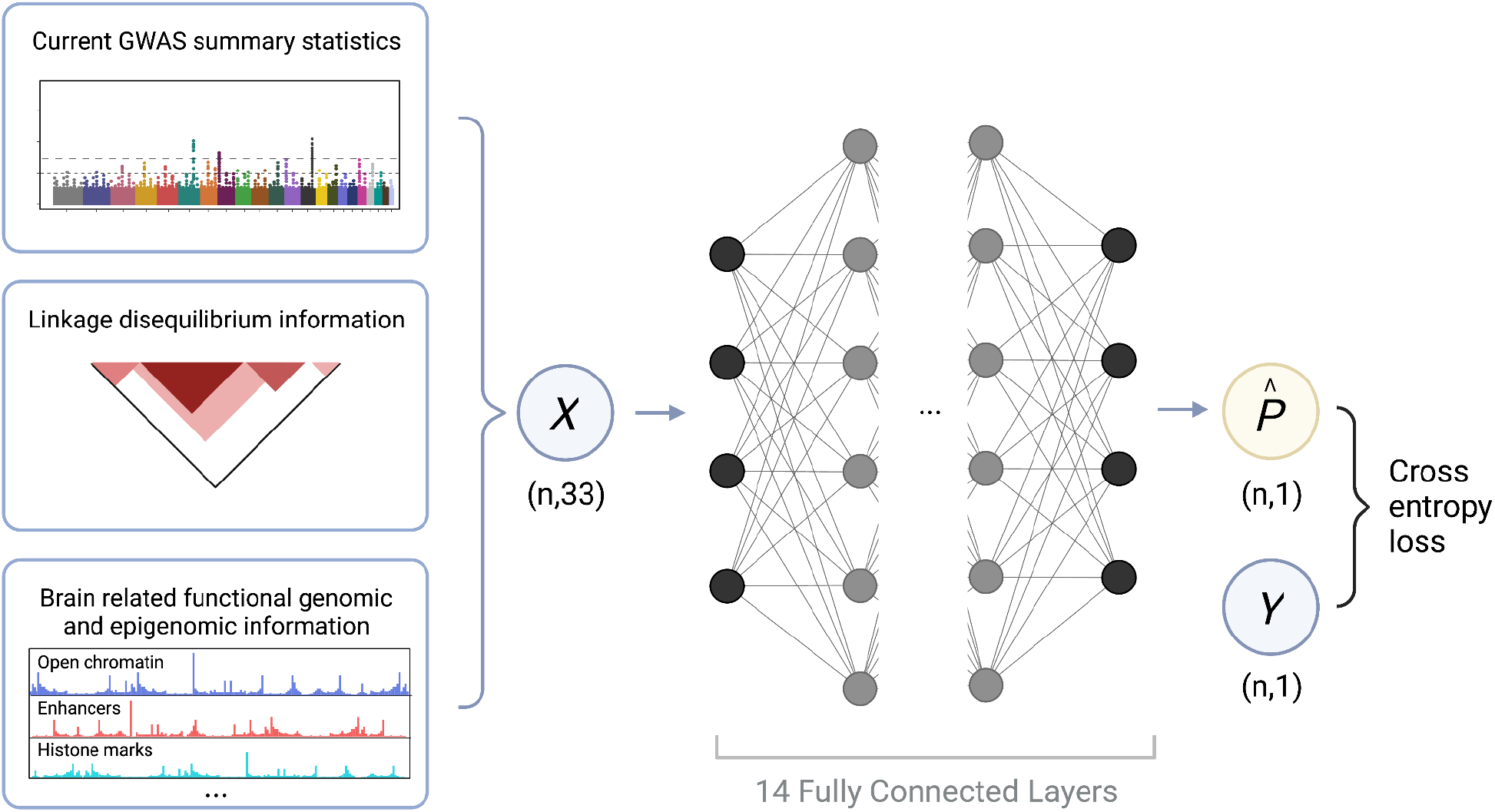
Overview of the DeepGWAS model. The blue circle *X* denotes the 33 input features, which serve as predictors in DeepGWAS model; the blue circle *Y* denotes the true binary input label indicating whether a variant is associated with the disease. During training, *Y* is obtained from a larger-sample-size study, serving as the working truth; *n* denotes the number of genetic variants; the black and gray solid circles denote the neural network nodes of the deep learning architecture within the DeepGWAS model; the yellow circle denotes DeepGWAS output 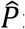 estimated probabilities for each of the *n* variants being associated with the disease.

**Fig. 2.**
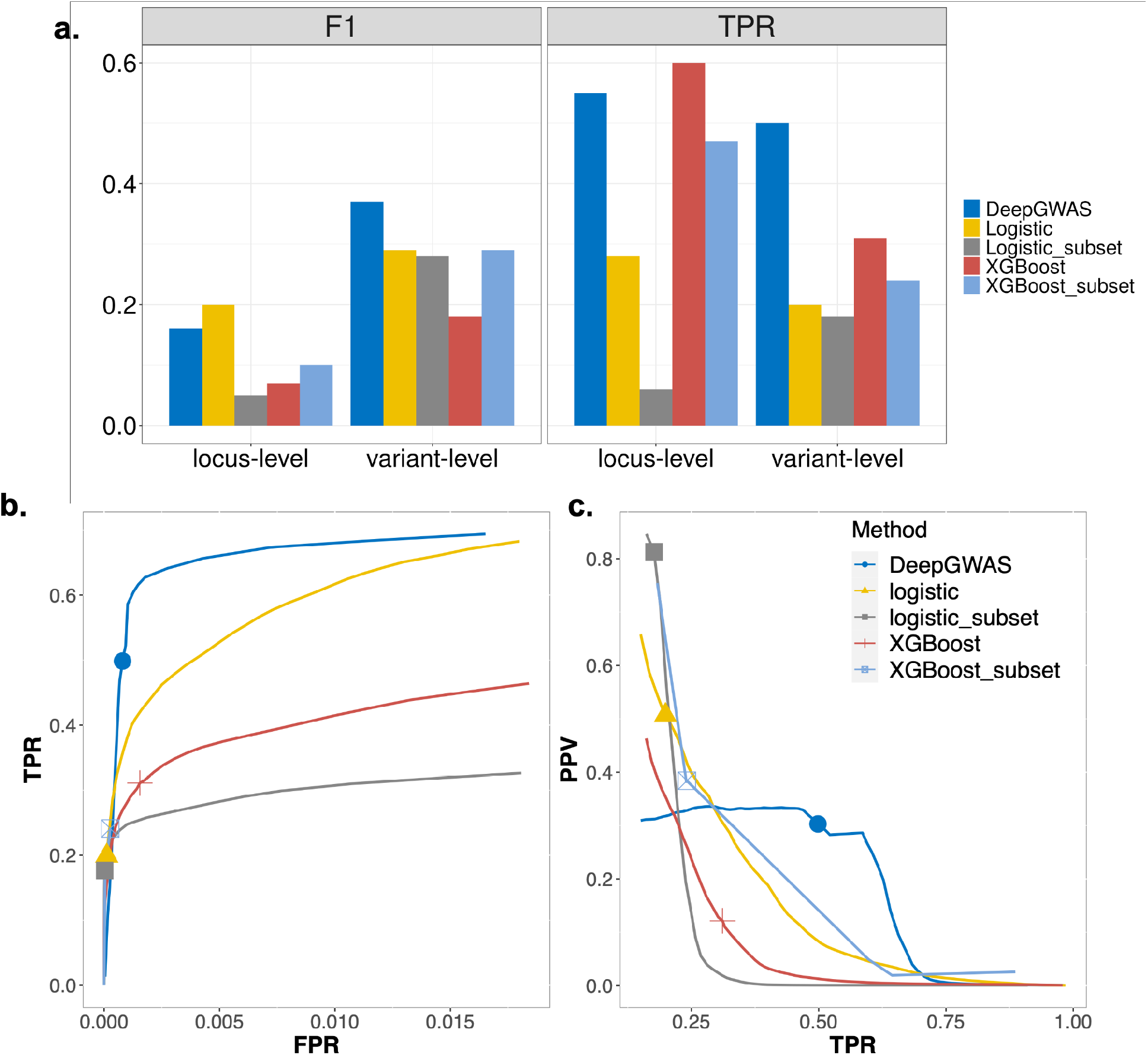
Model comparison in SCZ data. **a**, Comparison of TPR and F1 at both variant and locus level for 5 methods; **b**, ROC curve for comparison at variant-level of logistic regression, XGBoost and DeepGWAS models for evaluation. Logistic is the logistic regression model based on all variants, while logistic_subset uses the same variants selected by DeepGWAS’ insignificant variant under-sampling strategy. Similarly for XGBoost and XGBoost_subset; **c**, Precision recall curve for comparison at variant-level of logistic regression, XGBoost and DeepGWAS models for evaluation. PPV: positive predictive value. Marked points in the lines denote results using prediction probability threshold value of 0.5. Abbreviations: FPR=false positive rate; TPR=true positive rate; ROC=receiver operating characteristic curve; F1=TP/(TP+½(FP+FN)).

### DeepGWAS enhancement reveals hundreds of novel SCZ loci

After systematically comparing DeepGWAS with alternative methods using 64 SCZ GWAS cohorts, we trained a DeepGWAS model using data from two recent European SCZ GWA studies, applied to the latest European SCZ GWAS, and investigated the enhancement results. Specifically, we trained a DeepGWAS model using GWAS summary statistics from Ripke et al. 2014, the 2nd largest European ancestry SCZ GWAS meta-analysis^4^, as input features, and using genome-wide significance (*p*-value < 5e-8) from Pardiñas et al. 2018 (the largest European SCZ GWAS)^18^, as the input *Y* label. Once trained, the DeepGWAS model was applied to the GWAS summary statistics from Pardiñas et al. 2018, and the results show that DeepGWAS, with the default threshold of 0.5, enhanced 413 loci compared to the input GWA study from Pardiñas et al. 2018^18^. Importantly, 88 out of 413 were validated by Trubetskoy et al. 2022, the most recent and largest multi-ethnic SCZ GWA study^5^ (**Fig. 3**).

**Fig. 3.**
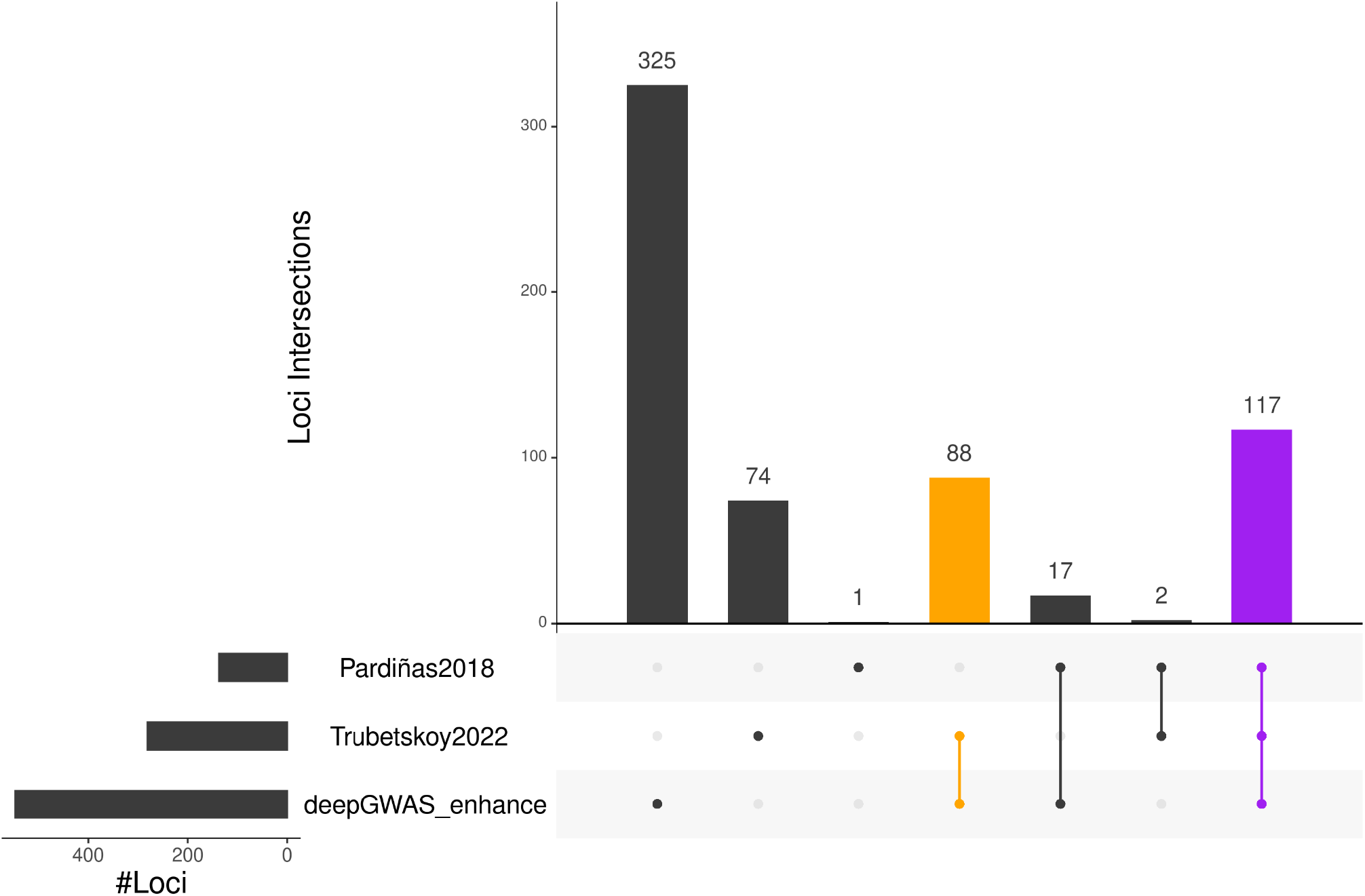
DeepGWAS enhanced Pardiñas et al. 2018 SCZ GWAS. Significant loci detected by each method/study are shown with an upset plot. The orange bar represents loci not in the input Pardiñas et al. 2018 GWAS results, enhanced by DeepGWAS, and validated by Trubetskoy et al. 2022; the purple bar corresponds to lower hanging fruit loci detected by all methods/studies, i.e., significant in the original input of Pardiñas et al. 2018, remain significant after DeepGWAS enhancement, and also significant in Trubetskoy et al. 2022.

### DeepGWAS’s enhancement via transfer learning suggests novel loci for AD and MDD

We have shown above that the DeepGWAS model trained using SCZ data has demonstrated satisfactory performance in enhancing SCZ GWAS. Importantly, we found that DeepGWAS could transfer the knowledge learned from SCZ data to enhance GWAS for additional neuropsychiatric disorders including Alzheimer’s disease (AD) and major depressive disorder (MDD) (“**Transfer learning using deepGWAS**”, **Online Methods**). Specifically, we fixed the model parameters for DeepGWAS learned from two recent European SCZ GWAS data described above, and applied the pre-trained DeepGWAS model to AD and MDD GWAS results.

### Enhancing AD GWAS via transfer learning

We applied the SCZ-trained DeepGWAS model to three AD GWA studies: Jansen et al. 2019^19^, Kunkle et al. 2019^20^, and Schwartzentruber et al. 2021^21^. There are three other AD GWA studies: Lambert et al. 2013^22^, Wightman et al. 2021^23^, and Bellenguez et al. 2022^24^. When applying the DeepGWAS model to each study, we used five other published AD GWAS to validate loci enhanced by DeepGWAS. From **Fig. 4** and **Fig. S1a-c**, 30% – 40% enhanced loci can be validated by other AD GWA studies, while few loci that were significant in the original input GWAS were missed by DeepGWAS. Taking the enhanced results based on Jansen et al. 2019^19^ as an example, we observe that DeepGWAS identified the *APP* locus which was not identified as a significant locus in the original Jansen et al., but was detected as a GWAS locus by several larger AD studies recently published^21,23,24^ (**Fig. 5**). *APP* is a well-established AD gene and previous studies have reported that mutations in *APP* can lead to *β*–amyloid protein accumulation and early-onset AD^25,26^.

**Fig. 4.**
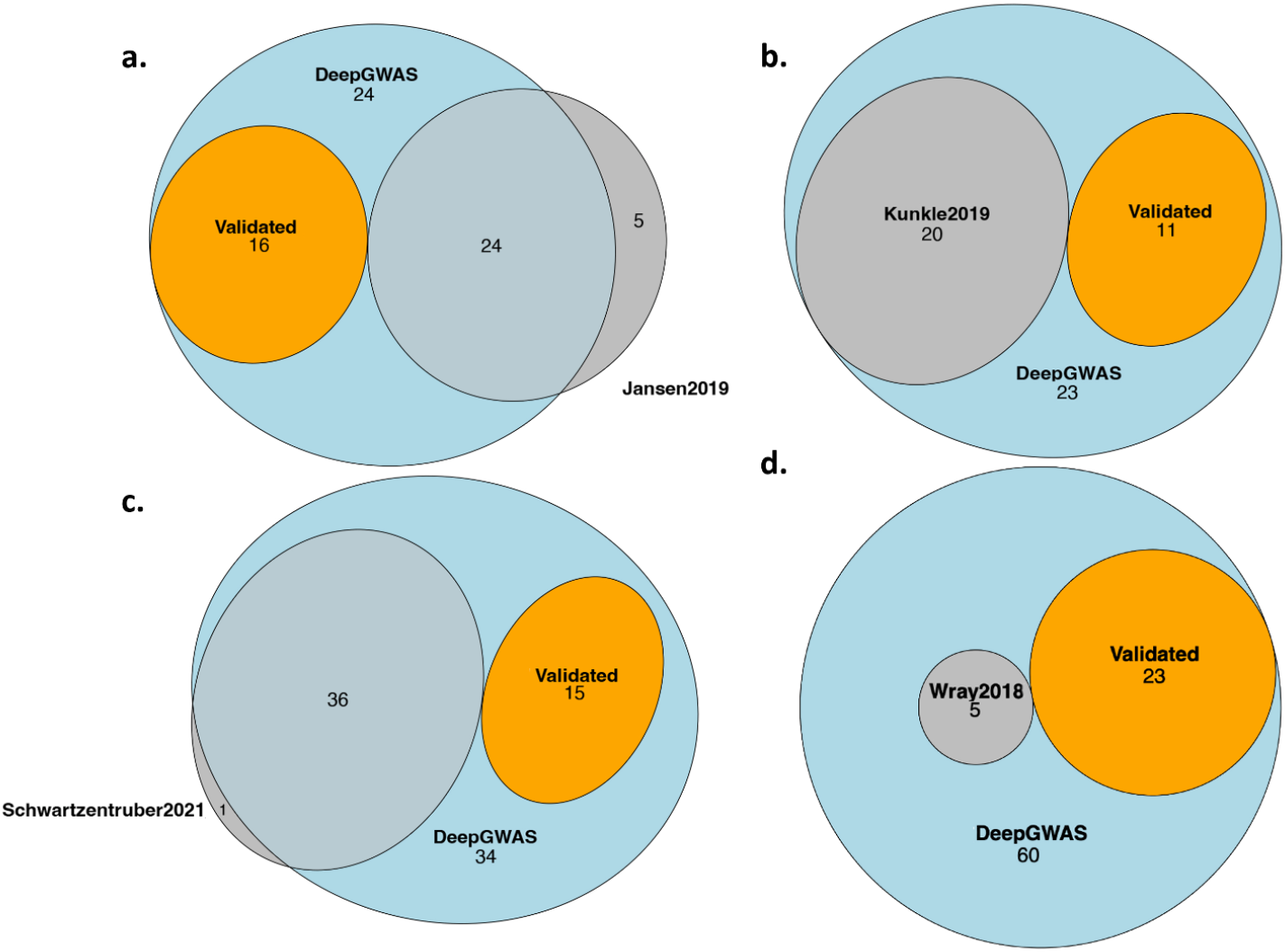
DeepGWAS performance when transferred to AD and MDD GWAS. **a**, Enhanced AD results when applied to Jansen et al. 2019^19^; **b**, Enhanced AD results when applied to Kunkle et al. 2019^20^; **c**, Enhanced AD results when applied to Schwartzentruber et al. 2021^21^; **d**, Enhanced MDD results when applied to Wray et al. 2018 (excluding 23andMe results)^29^. “Validated_loci” denotes DeepGWAS enhanced loci that can be validated by GWAS other than the input study.

**Fig. 5.**
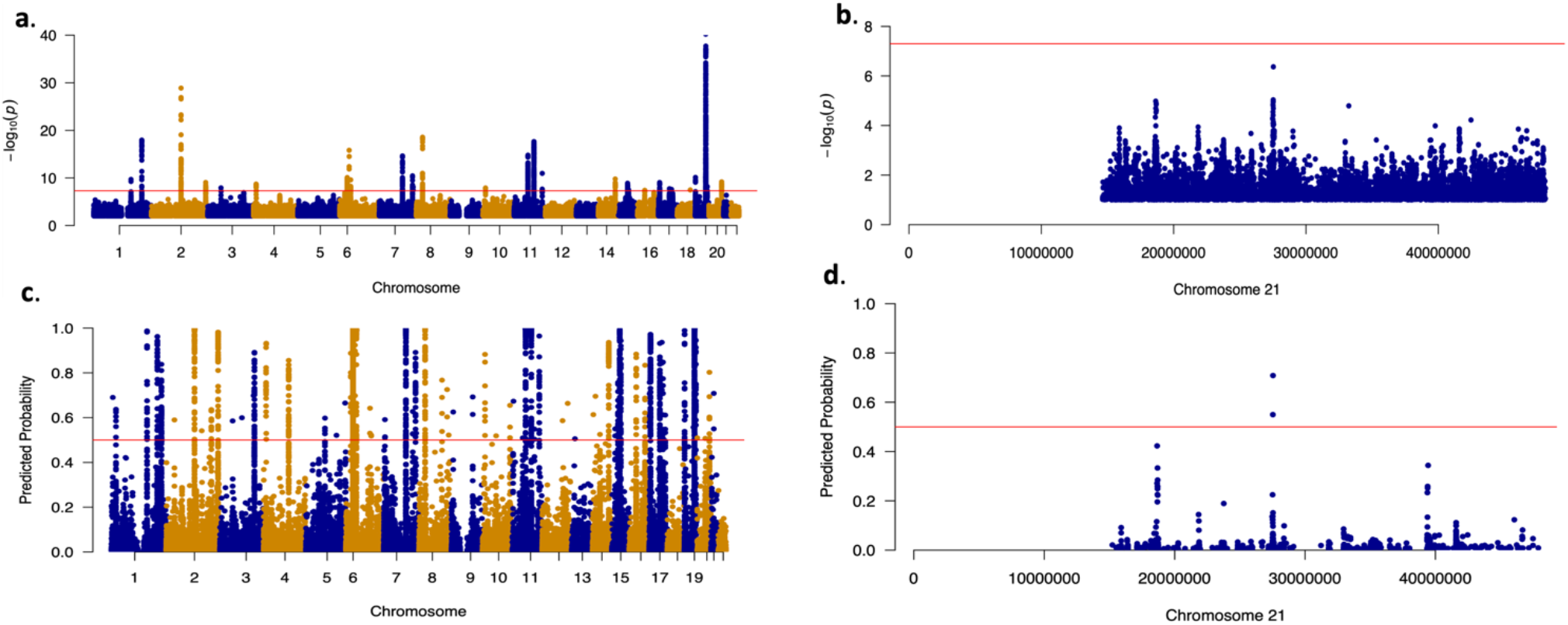
The *APP* locus enhanced by DeepGWAS. **a**, Manhattan plot to show the original GWAS results from Jansen et al. 2019^19^, and **b**, zoom in from **a** for chr21; **c**, Manhattan plot to show enhanced results by DeepGWAS. Y-axis is the prediction probability from DeepGWAS, and the red horizontal line marks 0.5. **d**, zoom in from **c** for chr21.

The *APP* gene serves as a positive control validated by earlier rare variant studies and more recent larger GWAS. In addition to *APP*, DeepGWAS identified other genes not, or not yet, validated by independent AD GWAS. Multiple genes have also been reported to be relevant to AD. For example, the locus marked by *EIF4G3*, the closest gene to a DeepGWAS index variant (i.e., the variant with the highest DeepGWAS predicted association probability at the locus) when applied to Jansen et al. 2019^19^, was reported as an AD locus in Naj et al. (2022)^27^. As another example, *NDUFAF6*, the closest gene to another DeepGWAS index variant, when the DeepGWAS model was applied to Kunkle et al. (2019)^20^ and Schwartzentruber et al. (2021)^21^, was reported to be associated with AD in a previous study using gene-wide analysis^28^. **Fig. S1d** summarizes DeepGWAS enhanced AD loci and the number of validated studies. Variants with higher DeepGWAS predicted probability are more likely to be validated by a larger number of studies. Interestingly and equally important, DeepGWAS also seems to be able to rectify potential false positives from standard GWAS. For example, one locus on chr18:56.18 MB, was significant in Jansen et al. 2019 but became non-significant after DeepGWAS. This anti-enhancement finding was confirmed by non-significance in three larger AD GWAS (Wightman et al. 2021^23^, Schwartzentruber et al. 2021^21^ and Bellenguez et al. 2022^24^.

### Enhancing MDD GWAS via transfer learning

We applied SCZ-trained DeepGWAS model to a MDD GWAS Wray et al. (2018)^29^ (excluding 23andMe due to their policies). We evaluated DeepGWAS enhanced results using a more recent MDD GWAS from Howard et al. 2019^30^, as well as Wray et al. 2018 full results including results from 23andMe. Results show that 22 out of 83 (∼26.5%) enhanced loci can be validated (**Fig. 4d, 6**), further demonstrating the transferability of the DeepGWAS model. For example, *KLF7*, the closest gene to a DeepGWAS index variant, when the DeepGWAS model was applied to Wray et al. (2018)^29^, was reported to be within a new MDD GWAS locus in Howard et al. 2019^30^.

*KLF7* as the target gene is further supported by adult cortex Hi-C data where the region harboring the DeepGWAS index variant (rs6717413) forms a significant chromatin loop with the promoter region of *KLF7*^31^ (**Fig. S2**). We note that there are 57 loci reported only by Howard et al. 2019^30^ (**Fig. 6**). These 57 loci remained non-significant even after DeepGWAS enhancement was applied on Wray et al. 2018 results, suggesting that increasing sample size is a power enhancer complementary to DeepGWAS’s computational enhancement for identifying additional variant-disease associations.

**Fig. 6.**
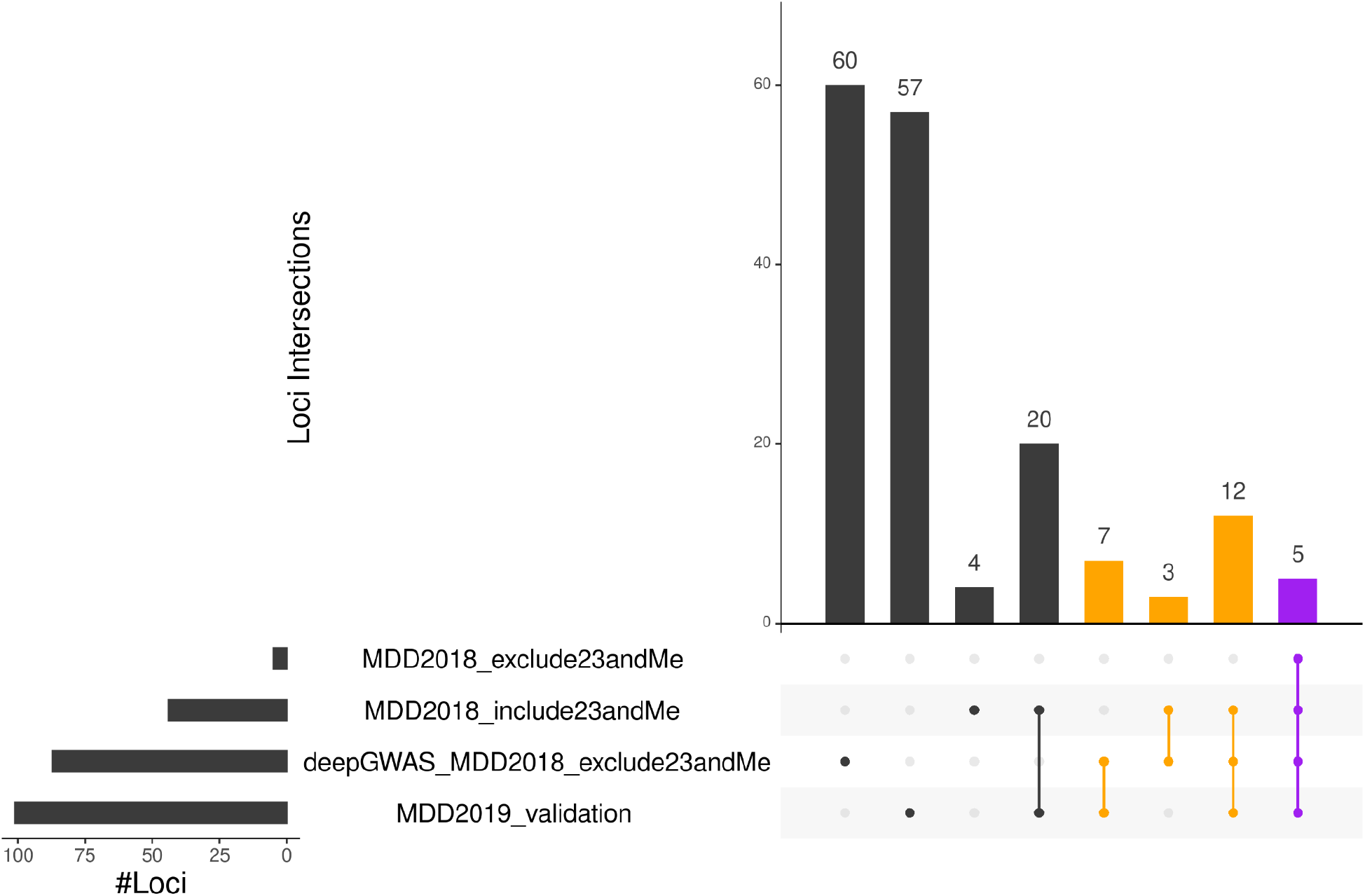
DeepGWAS results when transferred to MDD. The results are visualized by this upset plot, where the orange bars denote the validated loci which means the enhanced loci by DeepGWAS and validated by other studies; and the purple bar denotes the common loci identified by DeepGWAS, Wray et al. (2018) and Howard et al. (2019).

## Discussion

We proposed here a novel deep neural network to enhance GWAS signals without increasing sample size for neuropsychiatric disorders. Systematic evaluation using SCZ GWAS data and real GWAS enhancement showed DeepGWAS achieved the best performance compared to other two state-of-the-art deep learning methods.

Although DeepGWAS and fine-mapping methods are similar in terms of prioritizing variants, they are different in at least two aspects. First, DeepGWAS allows more complex relationships between loci and disease phenotype including non-linear relationships by employing a deep learning model. Second, DeepGWAS naturally accommodates both qualitative and quantitative annotations, while most fine-mapping methods only allow qualitative annotations. Among 33 features of interest, initial GWAS *p*-value is the most important feature, followed by super FIRE in adult annotations and LD score for known GWAS variants and eQTLs (**Fig. S3**). Specifically, approximately 69% ∼ 87% enhanced variants have initial *p*-values < 1e-5 in the input GWAS. In addition, DeepGWAS enhanced variants are more likely to reside in super FIRE regions, exhibit higher LD score for known GWAS variants, and are more likely to be eQTLs (details in “**Feature importance”** in **Supplementary Materials** and **Fig. S4**).

By default, we used DeepGWAS’s prediction probability 0.5 as the threshold, for screening purposes where our goal is to maximize power while tolerating false positives. Investigators may desire more stringent thresholds to shortlist variants or to reduce false positives. We investigated other threshold values from 0.5 to 0.95 with an increment of 0.05. We found 0.5-0.75 would be a good calibration threshold value interval and adapted to the user’s preferences based on our calibration on SCZ, AD and MDD studies (**Fig. S5a-e**). With higher or more stringent thresholds, DeepGWAS would detect fewer variants. For example, the number of DeepGWAS enhanced loci decreased from 413 to 39 for the testing SCZ GWAS dataset from Pardiñas et al. 2018 (**Fig. 3** and **Fig. S6**) when the threshold was increased from the default 0.5 to 0.9. Accordingly, the number of enhanced loci that can be validated by the independent Trubetskoy et al. 2022 decreased from 88 to 17. Similar trends were observed for MDD (**Fig. 6** and **Fig. S7**). Users therefore should choose thresholds that suit their purposes.

In this work, we trained our DeepGWAS model using two GWA studies on SCZ. Future efforts are warranted to train a “meta” DeepGWAS model with GWAS data from multiple genetically-correlated diseases, as different neuropsychiatric disorders like SCZ, MDD, and bipolar disorder are known to share some common genetic determinants as do certain neurological diseases (e.g., *APOE* in Alzheimer’s and Parkinson’s disease^32^. The immediate advantage of combining GWAS across diseases is to increase sample size for training, which in principle often improves the performance of neural network performance.

Importantly, DeepGWAS had the ability to transfer knowledge from one disease (SCZ) to other diseases (AD and MDD). DeepGWAS model can potentially transfer knowledge from one neuropsychiatric disorder to other neuropsychiatric disorders such as bipolar, autism, and Parkinson’s disease. It is also worthwhile to assess whether DeepGWAS model can transfer knowledge to other non-neuropsychiatric diseases or traits, because increasing sample size is generally expensive for GWAS of almost any trait. For disorders or traits not directly brain-related, annotation matching by tissue and cell type would be a non-trivial task that warrants separate future studies. Nevertheless, we believe our DeepGWAS model is a generalizable and valuable approach to enhance GWAS with additional knowledge that may be relevant to the diseases or traits under study. Careful training with SCZ data and applications to SCZ, AD and MDD GWAS presented in this work have demonstrated DeepGWAS’s enhanced power as well as the potential to remove false positives in the original study, by integrating GWAS results with relevant annotations in a deep learning framework.

## Online Methods

### DeepGWAS model

DeepGWAS is a fully connected deep neural network model, which aims to enhance GWAS results (**Fig. 1)**, by discovering additional candidate loci relevant for complex diseases or traits. The structure of DeepGWAS model utilizes 33-dimensional vectors as predictors (input), including GWAS summary statistics such as *p*-value and odds ratios as well as population genetics metrics such as MAF and two different LD scores (specifically, the regular LD score [summing across all variants] and LD score with significant variants in the input GWAS), which could also be calculated from a matching reference panel if no individual level data available.

MAF and LD scores calculation require a matching ancestry reference panel otherwise it may affect the performance of DeepGWAS model. While for the 28 annotation-related features including brain open chromatin regions and eQTLs, users can use the released annotations or complement more annotations and assemble those 28 categories to apply DeepGWAS model. The output of DeepGWAS is each variant’s predicted probability of being associated with the trait/disease of interest. We denote the DeepGWAS model as *F*, the input SNP feature matrices as *X*, the input binary label as *Y*, and the predicted probability 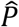 as F(*X*) (**Fig. 1**). Binary cross entropy loss is adopted in the training process. The goal of training is to learn *F* that minimizes the binary cross entropy (details “**DeepGWAS model**” in **Supplementary Materials**).

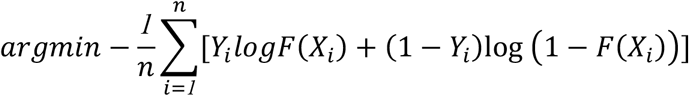

We aimed to release the pre-trained DeepGWAS model using the latest European SCZ GWAS summary statistics. To achieve the aim, we trained DeepGWAS model using Ripke et al. 2014, the 2nd largest European SCZ GWAS^4^ that identified 108 loci with 36,989 cases and 113,075 controls and using Pardiñas et al. 2018, the largest European SCZ GWAS^18^ that identified 145 loci with 40,675 cases and 64,643 controls (**Table S3**). Both sets of European GWAS summary statistics could be downloaded from https://www.med.unc.edu/pgc/download-results/. The SCZ-trained DeepGWAS model is released on Github https://github.com/GangLiTarheel/DeepGWAS.

### GWAS summary statistics

DeepGWAS performs analysis on GWAS summary statistics. In this work, we used the following summary statistics for SCZ, AD, and MDD GWAS.

#### Schizophrenia (SCZ) GWAS data

We assembled SCZ GWAS summary statistics from a total of 64 European cohorts, all contributing to the latest SCZ GWAS meta-analysis^5^. The sample sizes in each cohort range from 389 to 12,310, with 204 to 5,370 SCZ cases, and the number of pre-imputation variants released varies from 225,788 to 813,688. Detailed information of 64 cohorts is listed in **Table S1**. The released DeepGWAS model was trained using the two largest European SCZ GWAS summary statistics^4,18^ (**Table S3**).

#### Alzheimer’s disease (AD) GWAS data

Six most recent Alzheimer’s disease (AD) GWAS data were used in our study^19-24^ (**Table S3**), which identified 20∼75 loci with sample size from 54,162 to 1,126,563 for AD and/or proxy AD. Since we can only download restricted GWAS summary statistics when we performed DeepGWAS analysis due to provisions in the data-sharing agreement, we only applied SCZ-trained DeepGWAS model to 3 studies instead of all AD studies, specifically Jansen et al. 2019^19^, stage I summary statistics in Kunkle et al. 2019^20^, and Schwartzentruber et al. 2021^21^ separately, and then used the rest of the five AD studies to validate enhanced loci.

#### Major depressive disorder (MDD) GWAS data

Two most recent major depressive disorder (MDD) GWAS data were used in our study^29,30^ (**Table S3**), each identified 44 loci with sample size of 480,359 and 102 loci with sample size of

807,553. Since only GWAS summary statistics excluding 23andMe in Wray et al. 2018 and Howard et al. 2019 can be downloaded from (https://www.med.unc.edu/pgc/download-results/), we only applied the SCZ trained-DeepGWAS model to Wray et al. 2018 excluding 23andMe study, and then used Howard et al. 2019 to validate the enhanced loci.

### Input features

#### GWAS summary statistics and population genetics metrics

Basic predictors included GWAS summary statistics and population genetics metrics in the DeepGWAS model. GWAS summary statistics included -log10(*p*-value), odds ratio (OR); and population genetics metrics included minor allele frequency (MAF) and LD scores. MAF was extracted from European ancestry individuals in the 1000 Genomes Project^33^. We calculated two LD scores for each variant: overall LD score and LD score with known variants where the known variants were defined as the significant variants in standard GWAS analysis. The regular LD scores were calculated by summing up LD *r*2 between the target variant and all other variants located within 1Mb of each target variant based on European individuals from 1000 Genomes Project. Similarly, LD scores with known variants were calculated the same way but only summing over LD pairs involving significant variants in the input GWAS, again within 1Mb of each target variant.

#### Functional annotations

Functional annotations were collected from rich resources. We briefly introduce each annotation feature below and detailed information is summarized in **Table S4**.

**Brain eQTLs -** Brain-related eQTLs were collected from three sources including eQTLs from 13 Brain regions from GTEx v8^34^; brain eQTLs from PsychENCODE; and eQTLs from Qi et al. 2018^35^. We combined these eQTLs and filtered at nominal *p*-value < 1.0e-6.

**Pathogenic annotation** - Pathogenic annotations included phyloP scores derived from vertebrate mammals model^36-38^, Fathmm-XF score^39^, and CADD-phred score^40^.

**Open chromatin regions -** Open chromatin regions were taken union of the open chromatin regions for adult and fetal from several published studies^12,41-44^.

**FIREs and super FIREs -** Frequently Interacting Regions (FIREs, 40KB resolution) and super FIREs were downloaded from^45^.

**Selective sweep regions** - We also collected selective sweep regions in European detected in 1000 Genomes Project using S/HiC^46,47^.

**ENCODE3 cCREs -** The candidate cis-regulatory elements (cCREs) from ENCODE3 were collected based on DHS (DNase I-hypersensitive sites), H3K27ac, H3K4me3, CTCF and transcription factors (TF)^48,49^ and were downloaded from https://www.vierstra.org/resources/dgf. **Additional epigenomic annotations** - We also collected 30 additional epigenomic annotations from ^12^. Since we have multiple similar open chromatin and histone features collected from ^12^, we adopted data-driven strategy to merge similar annotations and used the Jaccard similarity index (bedtools v2.29.0) to group them into 11 meta-annotations (details in “**Data-Driven Clustering for epigenetics annotations”** in **Supplementary Materials**) which was shown in **Fig. S8**, and merged the annotations within the one sub-annotations using bedtools (v2.29.0). With all the functional annotations above, we have in total 30 functional annotations used as predictors in the DeepGWAS model.

### Evaluation using SCZ GWAS results

With GWAS summary statistics from 64 SCZ studies, we first randomly split them into three sets: set A, set B and set C (**Fig. S9**). Each variant in set A, B and C was annotated for all features listed in the “***Functional annotations***” section above. When splitting, we made efforts to balance the three sets considering several aspects including the number of cases, total sample size (i.e., the number of cases and controls), and the number of significant variants. To mimic increasingly larger GWAS, we assigned 10, 22 and 32 studies to set A, B and C respectively. After splitting, we meta-analyzed GWAS within each set using METAL^50^, and obtained three sets of GWAS summary statistics (**Fig. S9)**. With these three sets of GWAS summary statistics, we first trained models to “enhance set A to set B”. In other words, set A contributed input features (*X* in **Fig. 1**) while set B contributed outcome labels (*Y* in **Fig. 1**). Specifically, the binary indicator of whether the meta-analysis *p*-value < 5e-8 from set B was used as *Y* to train models. We then applied this pre-trained model to set B, to obtain enhanced set B results. Finally, significance in set C was served as ground truth to evaluate enhanced set B results.

We repeated the splitting procedure, randomly generated a new independent testing data following the same evaluation procedure, and applied the pre-trained DeepGWAS, XGBoost, and logistic regression three models to another independent testing dataset. We finally calculated the mean of the F1 score, CPR, TPR, ROC metrics, and precision recall curve metrics. The detailed information including the sample sizes and loci number used in evaluation were included in the **Table S2**.

### Comparison with alternative methods

To evaluate the performance of the DeepGWAS model, we compared DeepGWAS with alternative methods including logistic regression and XGBoost.

#### Logistic regression model

We trained a logistic regression model implemented in R v3.6.0. Logistic regression model was formulated as below, β_!_ denoted as weights of predictors and *X*_!_ denoted predictors. The output of the logistic regression model was prediction probability 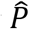 of whether a given variant be significantly associated with a disease.

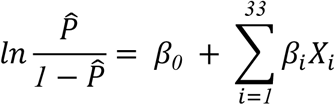

#### XGBoost model

XGBoost, or eXtreme Gradient Boosting, is a commonly used decision-tree-based ensemble machine learning algorithm using a gradient boosting framework. Using the same training dataset and testing dataset as applied to the DeepGWAS model, we trained and tested a supervised XGBoost model in R v3.6.0. We specified the learning task to be a tree-based logistic regression and evaluation metric to be root mean square log error (RMSLE). We set maximum boosting iteration as 50 in the model. Due to the extreme unbalanced ratio between the significant variants versus insignificant variants in the model, we used the argument of scale_pos_weight to control the unbalanced data.

To assess the performance of three models, we first defined the enhanced variants and loci as the predicted significant variants and loci that are not considered to be significantly associated with the disease in the input study. Then, we considered two metrics: truth positive rate (TPR), and F1 score. In addition, receiver operating characteristic curves (ROC) and precision recall curves (PRC) were also used to compare three models.

### Transfer learning using DeepGWAS

Although DeepGWAS training is supervised with labels from a large-sample-size study of the same disease (**Fig. 1 and Fig. S9**), DeepGWAS can transfer the knowledge learned from one disease (SCZ) to other diseases (such as AD and MDD). Specifically, we first trained our DeepGWAS model with two largest European SCZ GWAS^4,18^, fixed all the parameters in the neural networks, and applied the SCZ-trained-DeepGWAS model to enhance AD and MDD GWAS. Then we summarized the enhanced AD results by first binning them according to the probability of significant association with the disease 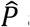and then assessing the proportion of loci within each bin that can be validated by independent AD GWAS (**Fig. S1d)**. The rationale behind this loci validation approach is that true positives are more likely to be enhanced by additional independent studies.

## Data availability

All GWAS data are available through the original publications with PMIDs listed in **Table S3**.

## Code Availability

This work uses the Plink v1.90.b3 software for LD clumping. Code for training and enhancing DeepGWAS model can be found at https://github.com/GangLiTarheel/DeepGWAS. We implemented and tested our DeepGWAS model under R 3.6.0 with keras_2.9.0 and tensorflow_2.9.0.

## Acknowledgements

We thank the Psychiatric Genomics Consortium for sharing their GWAS summary statistics. This project is funded by NIH grants R56-AG079291, U01DA052713, R01MH125236, and R01MH123724. J.W. and W.G. are supported by a grant from the National Institutes of Health, USA (NIH grant T32ES007018).

## Author Contributions

The study was designed by P.F.S and Y.L.. Funding was obtained P.F.S. and Y.L.. Analysis was performed by J.W., and G. L.. B.Q.L. and X.H. provided epigenomic annotations. J.W., G.L. and J.W.C. provided results visualization. Q.S. and W.F.L. provided analysis assistance. J.W., and G.L. wrote the manuscript. P.F.S, X.H. and Y.L. contributed to writing. All authors reviewed and approved the final version of the manuscript.

## Inclusions & Ethics

The procedures we followed were in accordance with the ethical standards of the responsible human rights committees on human experimentation.

## Additional information

Supplementary information including Supplementary Figs. 1–9 and Supplementary Note is available at xxxxx.

