## Supplementary Material and Figures for "DeepGWAS: Enhance GWAS Signals for Neuropsychiatric Disorders via Deep Neural Network"

### Supplementary Note

#### DeepGWAS model

DeepGWAS used 14 fully connected layers with 976 parameters to non-linearly map the patterns from 33 predictors to the predicted probability $\hat{P} = F(X)$. In each hidden layer, ReLU[1] is used as the activation except for the last layer with sigmoid function.

$$f_1(X) = max\ (0, w1 * X + b1)$$
$$f_2(X) = max(0, w2 * f_1(X) + b2)$$
$$\cdots$$
$$f_{33}(X) = max(0, w33 * f_{11}(X) + b33)$$
$$f_{34}(X) = \frac{1}{1 + e^{-(\ w34 * f_{33}(X) + b34)}}$$
$$\hat{P} = F(X) = f_{34}{}^\circ f_{33}{}^\circ \cdots {}^\circ f_1(X)$$

During the training process, we optimized $\{w1, w2, \ldots, w34, b1,\ b2, \ldots, b34\}$ to minimize the binary cross entropy loss in the training dataset. Backpropagation with Adam[2] was used as the updating strategy. In the current DeepGWAS model, we only considered the classic fully connected deep neural networks as the first deep learning work to solve this task. It might be of considerable interest to attempt other state-of-the-art deep neural network architectures, like convolution neural networks[3] and transformer neural networks[4], on the same GWAS prediction task. With the fast advancement of deep learning techniques, we expect the DeepGWAS model to further improve with more sophisticated architectures.

#### Under-sampling insignificant variants for model training

In a typical GWAS, only a small proportion of variants reach genome-wide significance[5,6]. Take the two SCZ GWAS summary statistics used for training data as an example, we have 7,548,429 variants (that are shared by both studies) in total, but only 10,802 (0.14%) variants were genome-wide significant[5] while 16,927 (0.22%) variants were genome-wide significant in the larger GWAS summary statistics designated to be the training dataset[6]. If we use all of the variants to train DeepGWAS, the model could easily be biased towards learning the class of variants not significantly associated with the disease since the majority of variants are insignificant. In practice, the model might converge to a trivial solution, which predicts all variants to be negative (i.e., not significant). To balance our training data, undersampling the negative class and oversampling the positive class are adopted. Additionally, we restricted the training data to consist of strictly positive ($p$-value < 1e-08) and strictly negative variants ($p$-value >0.1), since they would contain more accurate information for training. Finally, we obtained 57,386 variants for training, which consists of 17,386 strictly positive variants (i.e., twice of 8,693 original strictly positive variants in input GWAS before enhancement) and 30,000 strictly negative variants (randomly sampled from 6,144,707 strictly negative variants in current training data). Note, we point out that the ratio of positive to negative variants adopted in this model design is not a unique, optimal ratio required for achieving the best DeepGWAS model. In order to fairly compare the performance of the logistic regression and XGBoost models, we further sub-sampled the training dataset to have the same scale as that of the DeepGWAS model. Therefore, we in total have 5 models to compare, namely logistic, logistic-subset, XGBoost, XGBoost-subset and DeepGWAS.

**Locus definition**

To define loci in our **systematic evaluation**, we used a general method to that starts with identifying significant variants (i.e., enhanced results with prediction probability > 0.5 and association results with $p$-value < 5e-8), and defined each locus as a region where two consecutive variants were within 1Mb of each other. Each locus spanned the region from the first variant to the last variant in that locus. Independent loci were defined as non-overlapping regions with a >= 1Mb gap, where the gap between loci was defined as the genomic distance between the last variant of a locus and the first variant of the next locus.

For all published **GWAS** used in this study, we used the loci definition as defined by the corresponding publication. For the genome-wide enhanced results based on real GWAS, we used a similar strategy described in [7] to define loci. We first summarized the association results as the number of independent index variants. Index variants were defined as LD-independent and had $r2$ < 0.1 within the 3-Mb window and used prediction probability of 0.5 as the significant threshold value. Then we defined the associated clump including the left and right most variants with $r2$ < 0.1 with the index variant. We additionally added a 50Kb window to each side of the LD clumps and combined overlapping clumps into one single locus; meanwhile, we removed the locus only including one variant[7]. The reference used to define LD was based on European samples from the 1000 Genomes Project.

#### Feature importance

We assessed the feature importance via ablation study. We left one feature out of the DeepGWAS model at a time and re-trained DeepGWAS model using the rest of 32 features as input predictors with the same structure as our full DeepGWAS model. In order to quantify feature importance, we calculated F1 score with a prediction probability threshold value of 0.5, and compared it to the F1 score of the full DeepGWAS model.

#### Data-Driven Clustering for epigenetics annotations

We also collected 30 epigenomic annotations from [8]. It includes one DNaseq-seq annotations from fetal brain (FB1); one H3K27ac annotation from fetal brain (H3K27ac); three ATAC-seq measured chromatin accessibility data from brain organoid at three different time points (Organoid_0, Organoid_11, Organoid_30); four histone features including three H3K27ac ChIP-seq annotations from adult brain prefrontal cortex (PFC_H3k27ac), temporal cortex (TC_H3K27ac), and cerebellum (CBC_H3k27ac), one active enhancers from adult brain prefrontal cortex (PEC_Enhancers); one open chromatin regions defined by either ATAC-seq or DNase signals from adult brain prefrontal cortex (PEC_OCR); two ATAC-seq annotations from early human cortex in germinal zone (GZ) and cortical plate (CP); fourteen neuronal ATAC-seq annotations from adult brain (ACC_neuron, AMY_neuron, DLPFC_neuron, HIPP_neuron, INS_neuron, ITC_neuron, MDT_neuron, NAC_neuron, OFC_neuron, PMC_neuron, PUT_neuron, PVC_neuron, STC_neuron, VLPFC_neuron); and five cell-type specific ATAC-seq annotations from human iPS cells derived neuronal cells (CN, DN, NSC, GA)[8].

### Supplementary Figures

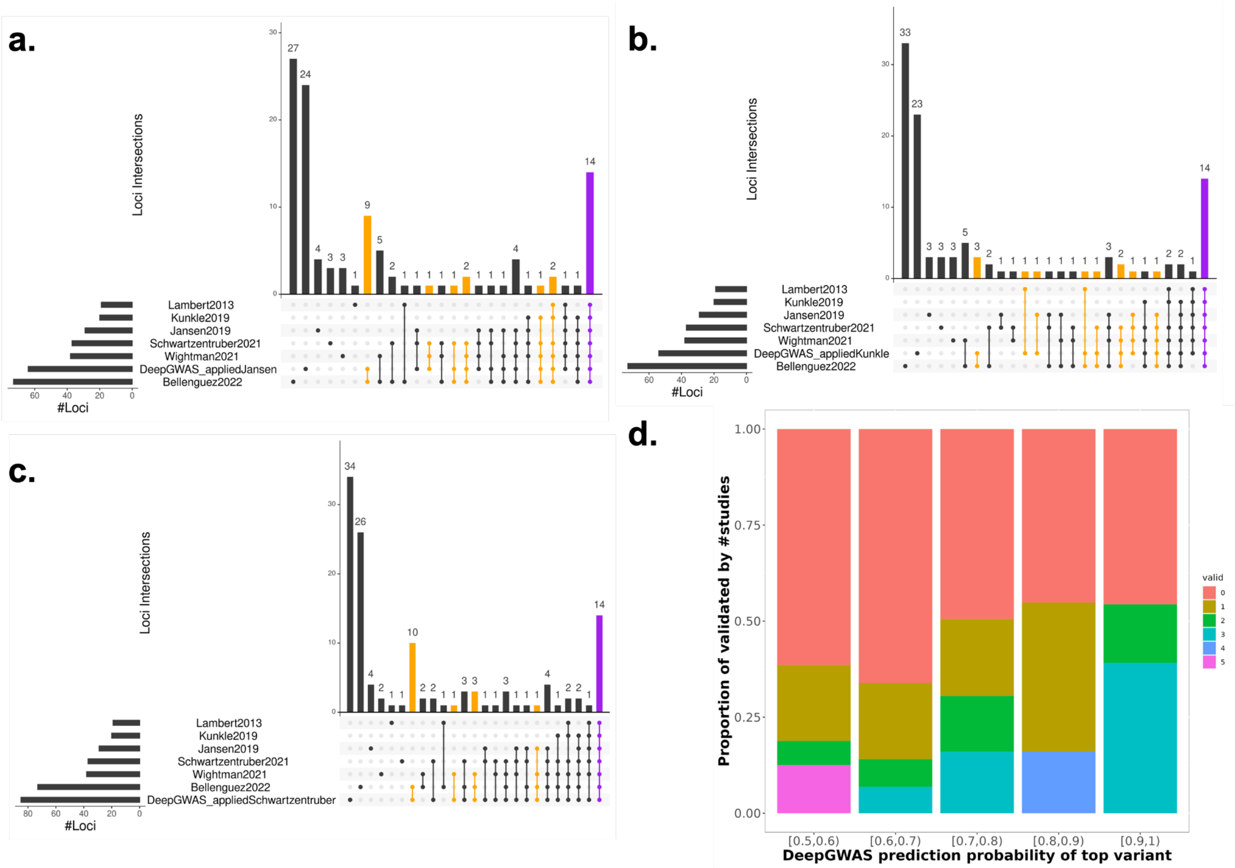


**Supplementary Figure 1. DeepGWAS enhanced results for AD.** Upset plots to show DeepGWAS results enhanced from input GWAS: **a**) Jansen et al. 2019[9], **b**) Kunkle et al. 2019[10], and **c**) Schwartzentruber et al. 2021[11]. The orange bars denote DeepGWAS enhanced loci that are also validated by other GWAS; the purple bar denotes the common loci identified by DeepGWAS and all AD GWA studies. **d**) Stacked bar plots to show the relationship between DeepGWAS confidence (measured by prediction probability, X-axis) and distribution of number of studies validating enhanced loci. For each bin of prediction probability (X-axis), we show proportions of enhanced variants (1) not yet validated by any other study (red, 0); (2) validated by one other published study (yellow, 1); (3) validated by two other published studies (green, 2); (4) validated by three other published studies (cyan, 3); (5) validated by four other published studies (blue 4); and (6) validated by all five other published studies (purple, 5).

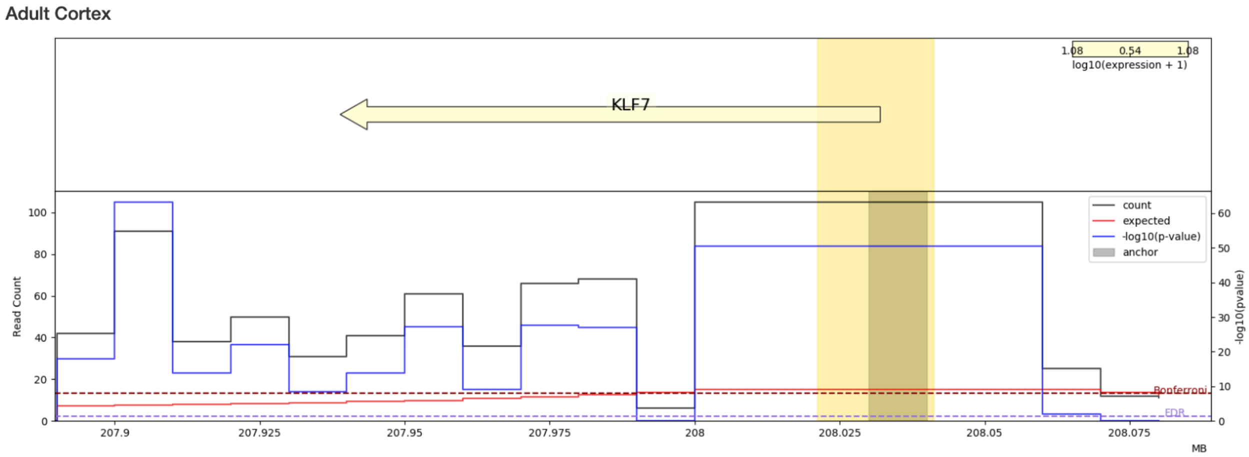


**Supplementary Figure 2. Virtual 4C plot for the index variant rs6717413 in adult cortex**. The bin containing rs6717413 is indicated by the thick gray vertical bar. The yellow vertical bar highlights the promoter region of *KLF7*. The black bars show observed Hi-C counts, red bars expected counts, and blue bars -log10(*p*-value) for chromatin interaction with the anchor bin (containing rs6717413). The range of -log10(*p*-value) is plotted on the right Y-axis while the range of the counts on the left Y-axis. The *X*-axis is genomic location in Mb.

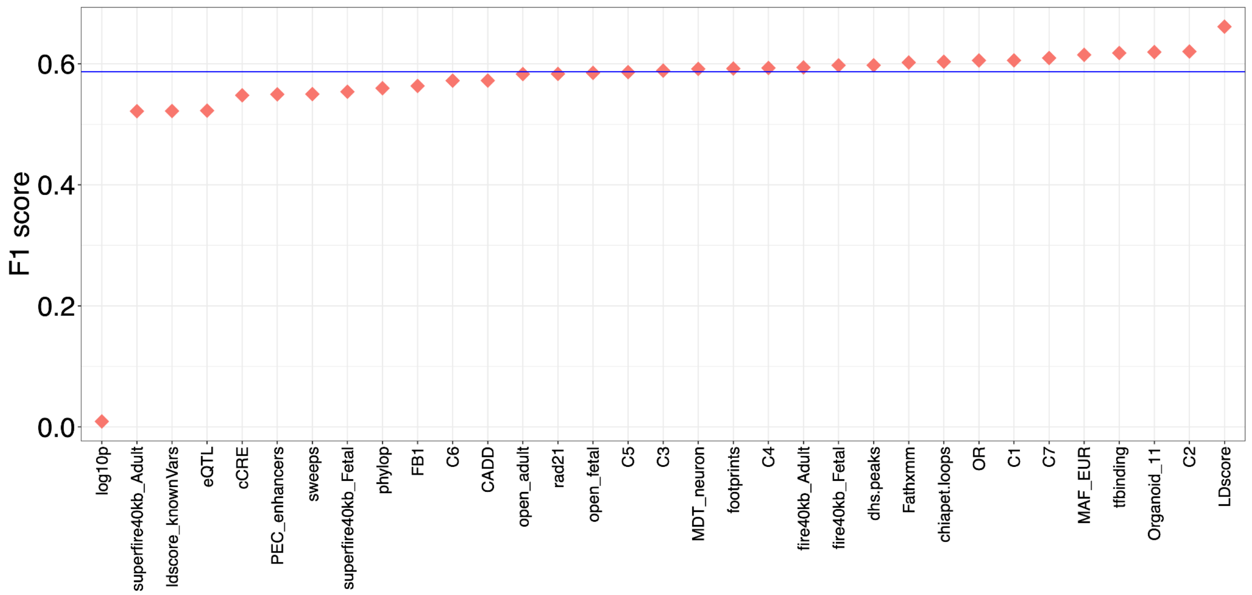


**Supplementary Figure 3. Feature importance evaluation from ablation study.** X-axis indicates the 33 models with each model removing one of the 33 features in the full/default DeepGWAS model. Y-axis shows the corresponding F1 score. The blue horizontal line denotes the F1 score from the full/default 33-feature DeepGWAS model.

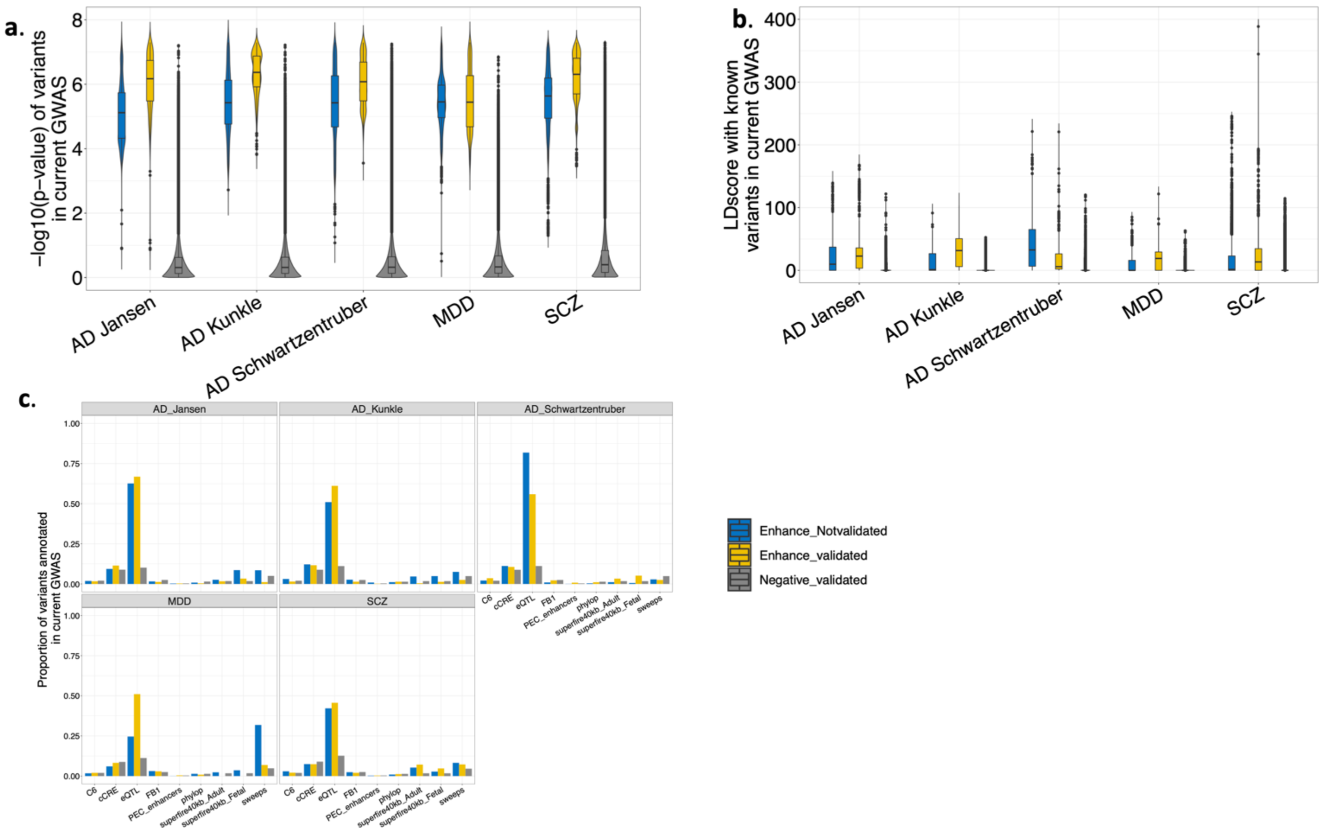


**Supplementary Figure 4. Characteristics of DeepGWAS enhanced variants.** Features used in DeepGWAS modeling are compared across three groups of variants: (1) "Enhance_Notvalidated" (blue): variants enhanced by DeepGWAS but not validated in other GWAS; (2) "Enhance_validated" (yellow): variants by enhanced DeepGWAS and validated in other GWAS; and (3) "Negative_validated" (grey): variants consistently insignificant across all GWAS (including the input GWAS and other GWAS) and pos-DeepGWAS enhancement. Violin plots comparing the distributions of **a**) -log10(*p*-value), **b**) LD score with known variants (i.e., significant variants in the input GWAS); **c**) bar plots showing the proportion of variants overlapping with annotation features (X axis). C6 denotes the open chromatin status, inferred from ATAC-seq data, from brain organoid. For details of other annotations, see "***Functional annotations***".

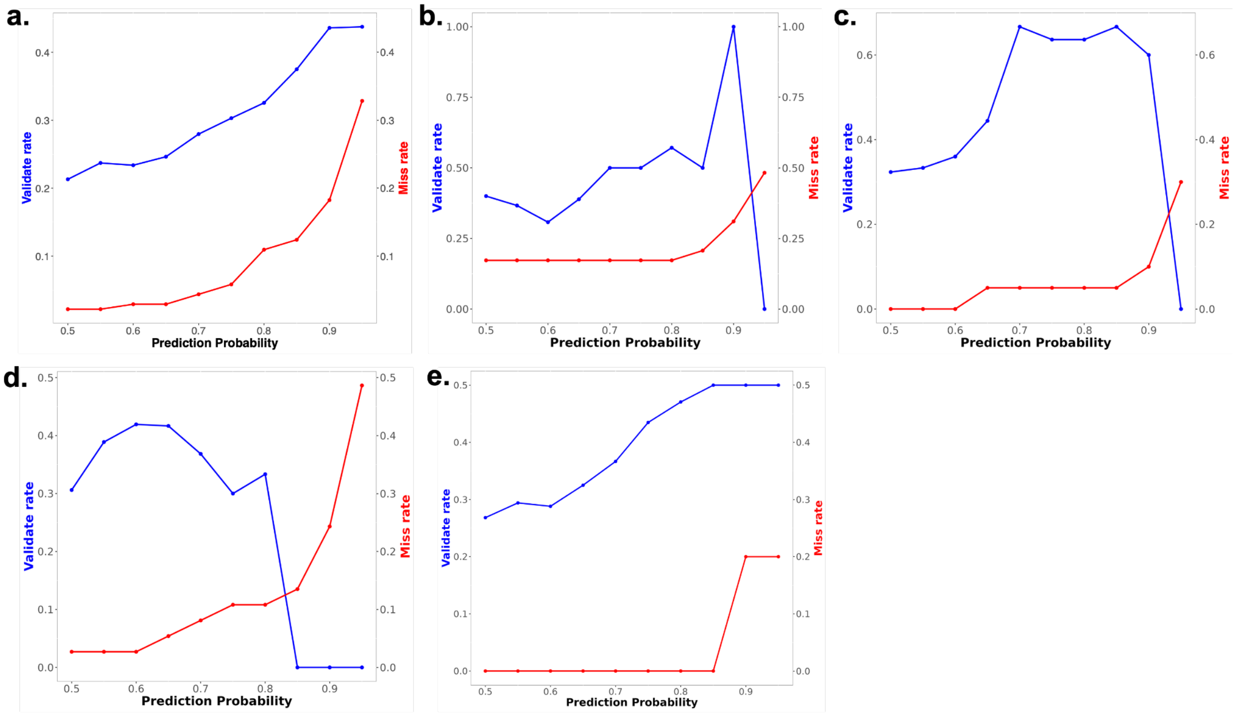


**Supplementary Figure 5. Performance under different thresholds of predictive probability.** Each of the 5 sub-figures shows the results of varying prediction probability thresholds for different input GWAS. **a**) Enhanced SCZ Pardiñas et al. (2018)[5]; **b**) Enhanced AD Jansen et al. (2019)[9]; **c**) Enhanced AD Kunkle et al. (2019)[10]; **d**) Enhanced AD Schwartzentruber et al. (2021)[11]; **e**) Enhanced MDD Wray et al. 2018[12]. The left Y-axis corresponds to the validation rate by the blue line and the right Y-axis corresponds to the missing rate by the red line.

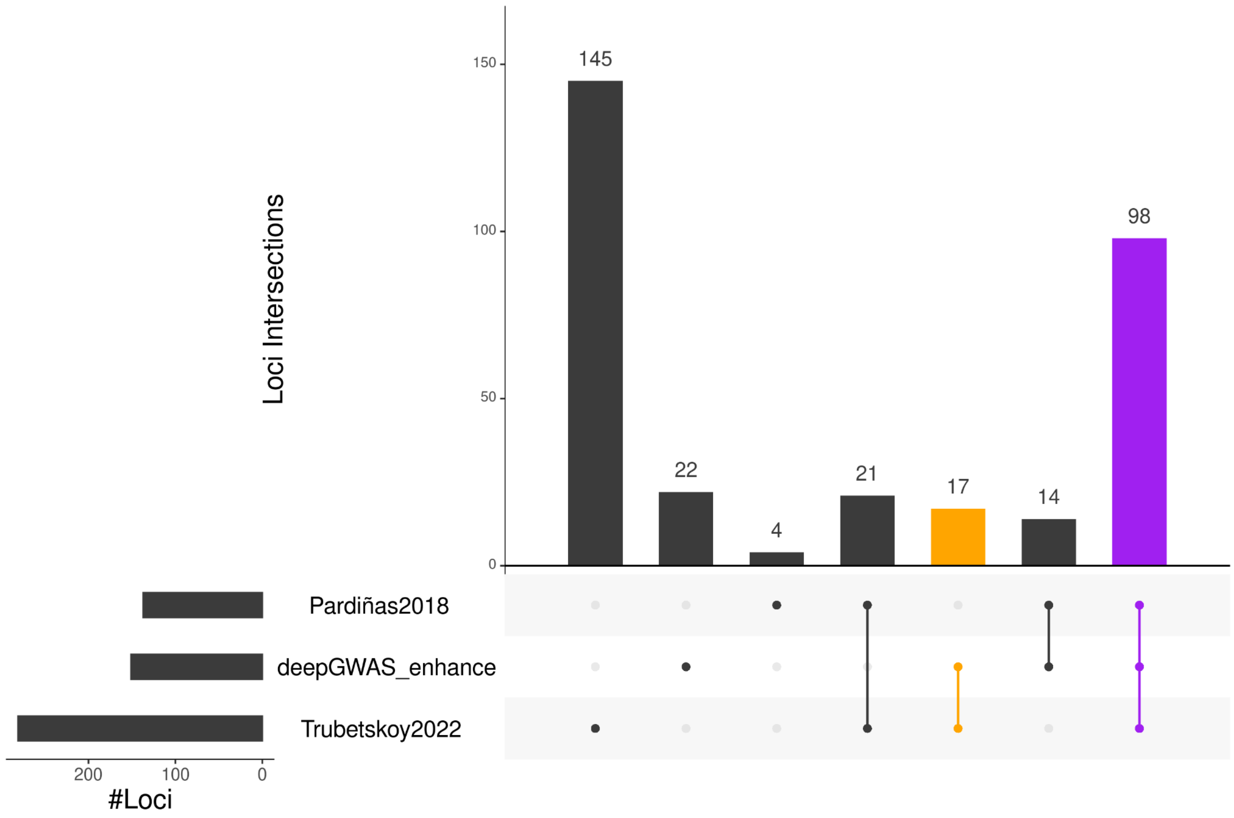


**Supplementary Figure 6. DeepGWAS enhanced Pardiñas et al. 2018 SCZ GWAS.** Significant loci detected by each method/study are shown with an upset plot. For DeepGWAS, the prediction probability threshold is 0.9. The orange bar represents loci not in the input Pardiñas et al. 2018 GWAS results, enhanced by DeepGWAS, and validated by Trubetskoy et al. 2022; the purple bar corresponds to lower hanging fruit loci detected by all methods/studies, i.e., significant in the original input of Pardiñas et al. 2018, remain significant after DeepGWAS enhancement, and also significant in Trubetskoy et al. 2022.

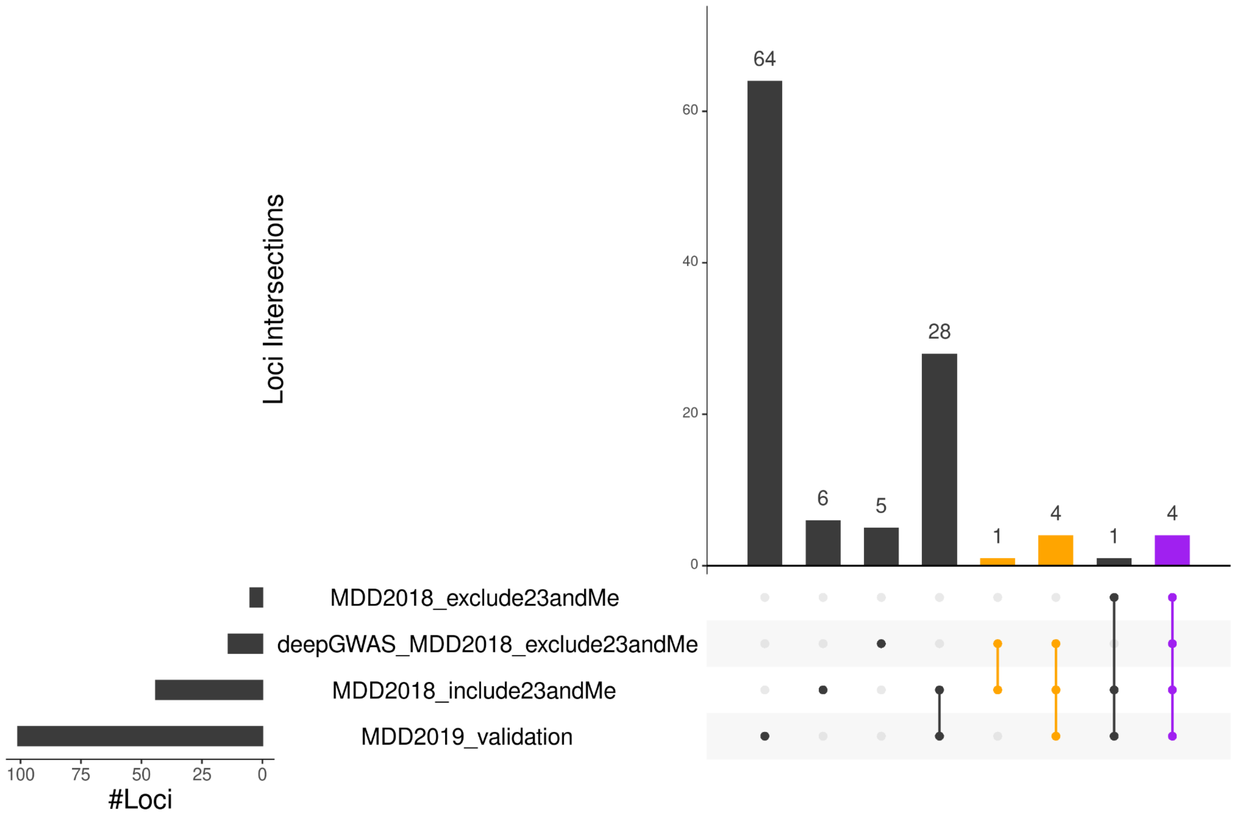


**Supplementary Figure 7. DeepGWAS results when transferred to MDD.** DeepGWAS results (with prediction probability threshold of 0.9), along with GWAS results, are visualized by this upset plot. The orange bars denote the validated loci which means the enhanced loci by DeepGWAS and validated by other studies; and the purple bar denotes the common loci identified by DeepGWAS, Wray et al. (2018) and Howard et al. (2019).

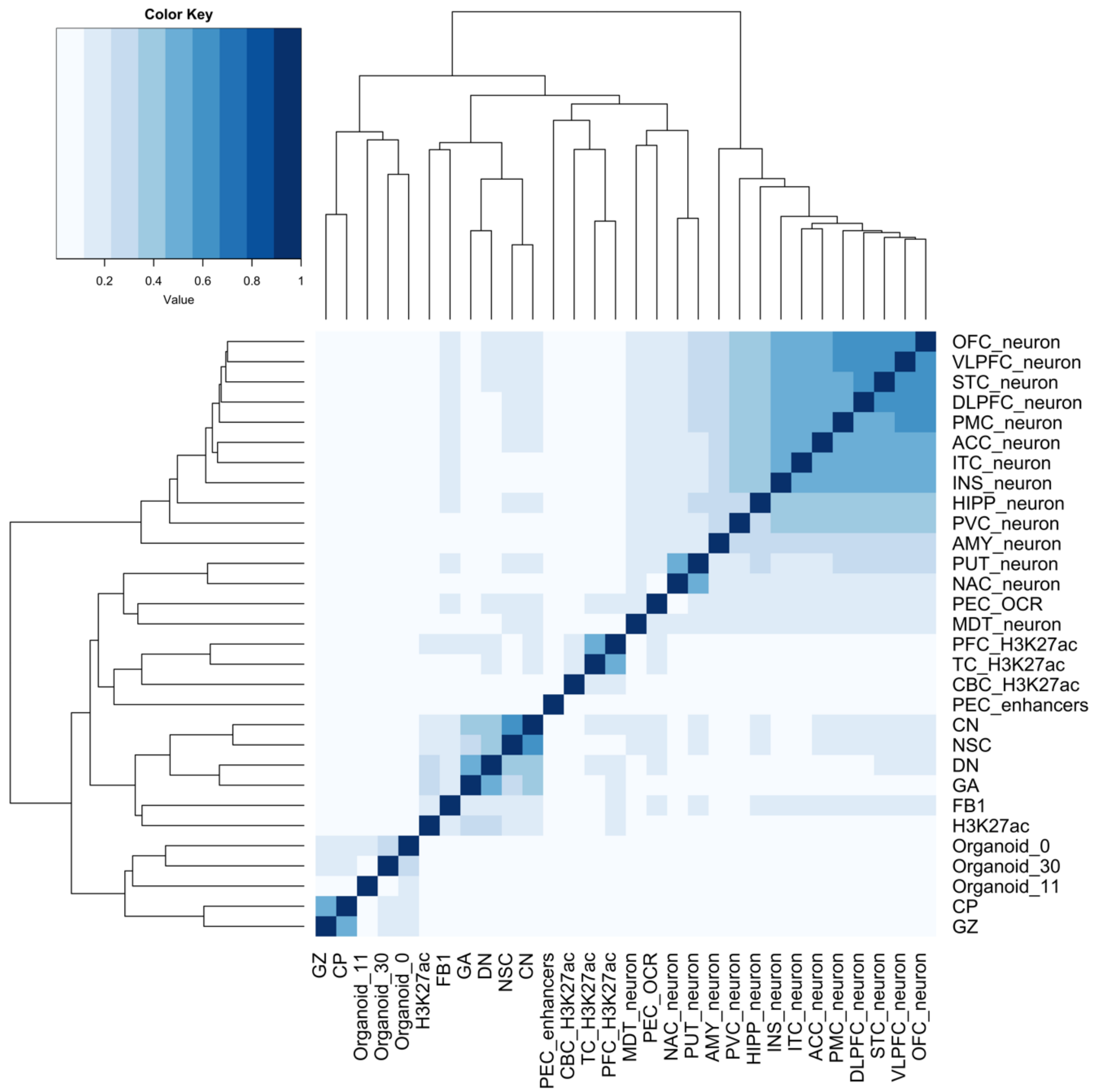


**Supplementary Figure 8. Jaccard indices for 30 epigenomic annotations**. Jaccard index is used to measure the similarity of any pair of epigenomic annotations. A higher value (darker blue in heatmap) indicates higher consistency or overlap. The dendrogram shows clustering results where similar annotations are grouped.

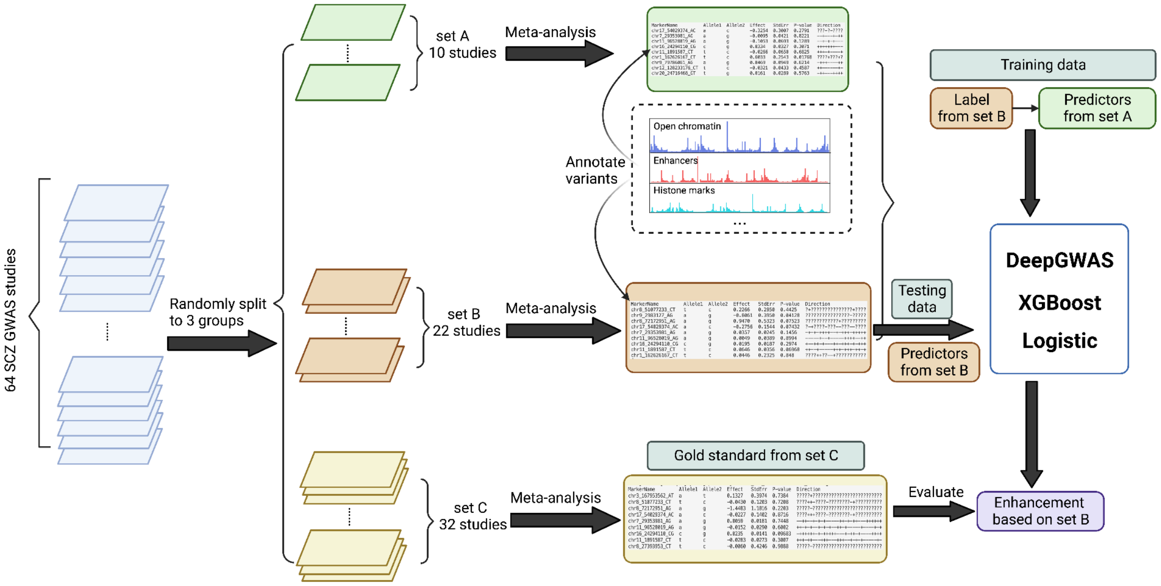


**Supplementary Figure 9. Construct DeepGWAS model using SCZ GWAS results.** GWAS results from 64 SCZ studies were randomly assigned to three sets (set A, set B and set C), and meta-analyzed within each set. We first trained our DeepGWAS, XGBoost and logistic regression models using predictors from set A and labels from set B. Then we used set B as the testing dataset where we applied the pre-trained DeepGWAS model on set B (i.e., where set B only contributed predictors) to predict enhanced labels in set B. Finally, labels from set C served as the gold-standard truth for evaluating the predicted enhanced labels in set B.
